# Adiponectin Reverses β-Cell Damage and Impaired Insulin Secretion Induced by Obesity

**DOI:** 10.1101/2022.07.22.501128

**Authors:** Ana Cláudia Munhoz, Julian D. C. Serna, Eloisa Aparecida Vilas-Boas, Camille C. Caldeira da Silva, Tiago Goss dos Santos, Francielle C. Mosele, Sergio L. Felisbino, Vilma Regina Martins, Alicia J. Kowaltowski

**Author notes:** Present address: Departamento de Biologia Celular e do Desenvolvimento, Instituto de Ciências Biomédicas, Universidade de São Paulo, São Paulo, Brazil (Vilma Regina Martins). Authors contributed equally (Julian D. C. Serna, Eloisa Aparecida Vilas-Boas).

## Abstract

Obesity significantly decreases life expectancy and increases the incidence of age-related dysfunctions, including β-cell dysregulation leading to inadequate insulin secretion. Here, we show that diluted plasma from obese human donors acutely impairs β-cell integrity and insulin secretion relative to plasma from lean subjects. Similar results were observed with diluted sera from obese rats fed *ad libitum*, when compared to sera from lean, calorically-restricted, animals. The damaging effects of obese circulating factors on β-cells occurs in the absence of nutrient overload, and mechanistically involves mitochondrial dysfunction, limiting glucose-supported oxidative phosphorylation and ATP production. We demonstrate that increased levels of adiponectin, as found in lean plasma, are the protective characteristic preserving β-cell function; indeed, sera from adiponectin knockout mice limits β-cell metabolic fluxes relative to controls. Furthermore, oxidative phosphorylation and glucose-sensitive insulin secretion, which are completely abrogated in the absence of this hormone, are restored by the presence of adiponectin alone, surprisingly even in the absence of other serological components, for both the insulin-secreting INS1 cell line and primary islets. The addition of adiponectin to cells treated with plasma from obese donors completely restored β-cell functional integrity, indicating the lack of this hormone was causative of the dysfunction. Overall, our results demonstrate that low circulating adiponectin is a key damaging element for β-cells, and suggest strong therapeutic potential for the modulation of the adiponectin signaling pathway in the prevention of age-related β-cell dysfunction.

Graphical Abstract:
Incubation of β-cells with sera or plasma from obese rats and humans hampers mitochondrial oxidative phosphorylation and glucose-stimulated insulin secretion (GSIS) relative to sera and plasma from lean rats and humans. Adiponectin, found at elevated levels in lean subjects, supports β-cell function on its own, in the absence of sera, and also reverses the effects of obese plasma. Prepared using Biorender.com.

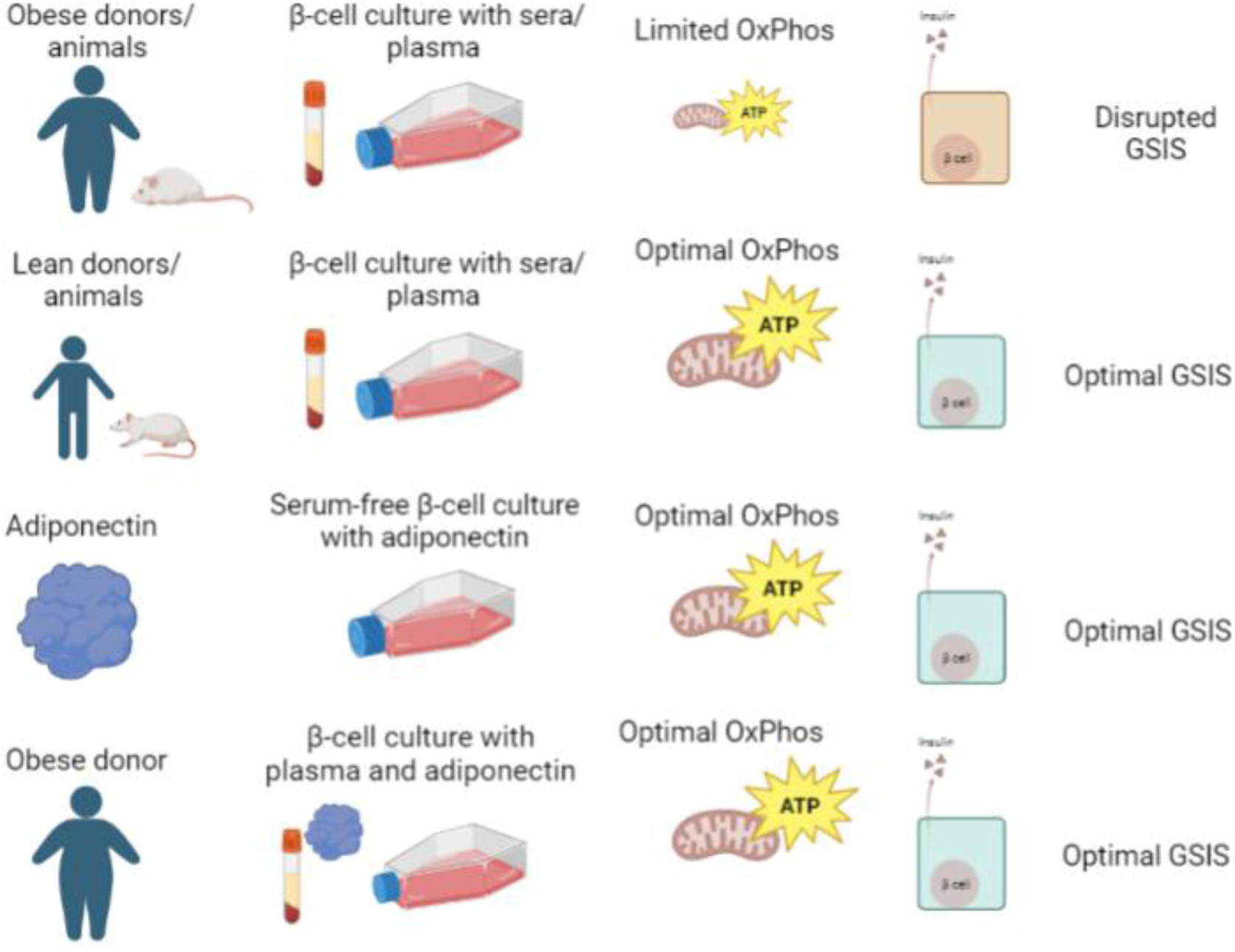

## Introduction

About half a billion persons currently live with type 2 diabetes mellitus, and this number is expected to increase by 51% until 2045 (Saeedi et al., 2019), as obesity indexes continue to rise and the population ages (Wang et al., 2011). Indeed, type 2 diabetes is a chronic age-related disease characterized by an inability to control blood glucose homeostasis, caused either by a decreased insulin action (insulin resistance) or inadequate insulin release by pancreatic β-cells (Kahn, 2003). Insulin resistance promoted by obesity has been related to changes in adipokines, inflammation, and is also affected by β-cell dysfunction, resulting in a loop mechanism (reviewed in Steppan et al., 2001; Kahn et al., 2006).

Pancreatic β-cells are susceptible to damage caused by excess circulating glucose and lipids (glucolipotoxicity), which is common in obese individuals (reviewed by Roma and Jonas, 2020). Indeed, a high calorie obesogenic diet, favoring glucolipotoxic conditions, is associated with type 2 diabetes and increased mortality (Hruby and Hu, 2015). On the other hand, caloric restriction (CR), preventing nutrient overload, is one of the most studied dietary interventions known to extend health and longevity, as well as prevent aging-associated β-cell dysfunction *in vivo* (Weindruch et al., 1986; Cerqueira et al., 2010; Hruby and Hu, 2015; Mattison et al., 2017; Hofer et al., 2022).

In a previous study (Cerqueira et al., 2016), we found that circulating factors in the sera from lean CR rats were highly protective for pancreatic islets, primary β-cells, and insulin-secreting cell lines *in vitro*, when added as a 10-fold diluted part of the incubation media, substituting commercial culture sera. Conversely, incubation for as little as 24 hours with sera from moderately obese animals fed *ad libitum* (AL) impaired insulin secretion. Sera from lean CR animals strongly protected β-cells from nutrient overload (glucolipotoxicity, Cerqueira et al., 2016), suggesting it contains soluble factors that enable enhanced insulin secretion under damaging conditions.

Glucolipotoxicity leads to β-cell dysfunction in a manner dependent on changes in mitochondrial structure and function (Molina et al., 2009). Indeed, mitochondria are central players in the loss of β-cell function related to type 2 diabetes (Las et al., 2020); in addition to their role regulating intracellular ATP production in response to glucose levels (and consequent insulin secretion), changes in mitochondrial genetics, bioenergetics, redox state, structure, and dynamics are involved in β-cell functional regulation (Las et al., 2020). During glucolipotoxicity, extensive mitochondrial fission occurs, promoting a fragmented and poorly interconnected mitochondrial phenotype and causing decreased mitochondrial ATP synthesis and β-cell dysfunction (Zhang et al., 2011; Cerqueira et al., 2016). Inhibiting this fragmentation prevents loss of cell viability, demonstrating that the change in mitochondrial morphology is causative of glucolipotoxicity-induced β-cell dysfunction (Molina et al., 2009). The protective effects of diluted CR sera on β-cells are also related to changes in mitochondrial dynamics and function, as it increases mitochondrial network interconnectivity and respiratory rates, without overt changes in mitochondrial mass (Cerqueira et al., 2016).

While previous work demonstrated the intracellular mechanisms in which CR sera promotes the preservation of β-cell function, the molecular factors from CR sera involved in this protection were not identified. The dilution of the sera as well as the fact it was collected from overnight-fasted animals and used in the presence of nutrient-rich culture media implies that changes in the amounts of substrates such as glucose, fatty acids, and amino acids are not involved in these results. Rather, the effects are most likely associated with hormones such as adipokines or other circulatory signaling components (Ouchi et al., 2011). Here, we investigated possible β-cell-protective factors in the sera of CR rats and lean humans, and identified adiponectin as a strong promoter of mitochondrial oxidative phosphorylation, insulin secretion, and β-cell preservation. Strikingly, adiponectin (Yamauchi et al., 2001) alone was able to promote β-cell function in the absence of any other serological component. It was also able to reverse β-cell dysfunction promoted by incubation with the serum or plasma from obese subjects, indicating a therapeutic potential for this pathway in the prevention of insulin secretion failure in obesity.

## Experimental procedures

### Animals, diets and serum collection

All experiments were approved by the animal use committees (*Comissão de Ética em Uso de Animais do Biotério de Produção e Experimentação da Faculdade de Ciências Farmacêuticas e Instituto de Química da USP* #109/18 and *Comissão de Ética no Uso de Animais do Instituto de Biociências de Botucatu,* UNESP, #1194).

Male 8-week-old Sprague Dawley rats were divided into two groups: AL, fed *ad libitum* with an AIN-93-M (Reeves et al., 1993) diet prepared by Rhoster (Campinas, Brazil) and CR rats, fed 60% of the AL group’s intake, using a diet supplemented with micronutrients to reach the same vitamin and mineral levels (Cerqueira et al., 2010). The animals were housed in groups of three per cage in 12 h light/dark cycles and given water *ad libitum*. The weight of the animals and food intake were recorded weekly to adjust the CR group diet to 60% of the intake of the AL group. At 34 weeks of age (after 26 weeks of the diet), rats were euthanized with a ketazine/xylazine overdose after 12 h fasting, blood was collected by cardiac puncture, and serum was obtained after clotting at room temperature for 30 min and centrifugation for 20 min at 300 x g. The supernatant was collected and stored at −20°C. Serum samples were thawed and heat-inactivated at 56°C for 30 min prior to use.

B6;129-Adipoq^tm1Chan^/J mice were obtained from The Jackson Laboratory (#008195, Bar Harbor, ME). Knockouts (KO) mice were backcrossed N11 to a C57BL/6J genetic background. Wild type C57BL/6J male 10-week-old mice were obtained from Institute of Biotechnology Animal Facility, Botucatu, Brazil. Both wild type and adiponectin KO mice were maintained in the Institute of Biosciences Animal Facility in cages on a ventilated shelf in 12 hours light/dark cycles, with *ad libitum* food and water. At 13 weeks of age, 12 wild type and adiponectin KO mice were euthanized, their blood collected by cardiac puncture, and serum was obtained after clotting at room temperature for 30 min and centrifugation for 20 min at 300 x g. The supernatant was collected and stored at −20°C. Serum samples were thawed and heat-inactivated at 56°C for 30 min prior to use.

### Pancreas histology

Whole pancreases from AL and CR animals were removed and immersed in 20 mL of 10% formalin for 24 hours. The tissue was then PBS-washed and dehydrated through a series of graded ethanol solutions (70, 80, 95 and 100%), followed by a clearing process with xylol, and finally embedded into paraffin blocks. The paraffin-embedded tissues were sectioned (5 µm thick) with a semi-automated rotary microtome (Leica Microsystems, Wetzlar, Germany) and placed on microscope glass slides coated with poly-L-lysine. Slides were stained with hematoxylin and eosin (H&E) and scanned using the TissueFAXS iPLUS (TissueGnostics, Wien, Austria) system under a magnification of 20. Three areas of each slide were blindly analyzed by two independent examiners using the ImageJ Fiji Software. Islets were categorized according to areas into very small (<1000 μm^2^), small (1000-5,000 μm^2^), medium (5,001-10,000 μm^2^), large (10,001-50,000 μm^2^) or very large (>50,000 μm^2^).

### Human plasma samples

Human plasma samples were obtained from the A.C. Camargo Cancer Center Biobank, and all experiments were carried out in accordance with the A. C. Camargo Cancer Center Institutional Review Board under registration n°. 3117/21. Samples include healthy donors who signed a free and informed consent form and authorized the institution to store and use their biological material for future studies. For the selection of research participants, data from medical records were examined and a balance in biological sex, ethnicity, age, eating habits, physical activity, and height was sought, separating subjects into lean and obese groups according to their body mass index (BMI). Subjects with pathologies, elevated blood pressure, smokers, regular alcohol consumption, sexually transmitted diseases, bariatric surgery, or chronic diseases were eliminated (Table 1). The overall selection includes patients of both sexes, equal in all quantifiable parameters except BMIs, producing four distinct groups: Lean women (BMI 22.0 ± 0.9, n=4), Obese women (BMI 31.0 ± 1.4, n=6), Lean men (BMI 23.3 ± 0.4, n=8) and Obese men (BMI 31.2 ± 1.4, n=10). Their blood was collected in a sterile vacuum tube containing 4.45 mmol/mL of the anticoagulant ethylenediaminetetraacetic acid dipotassium salt (EDTA). The blood was then centrifuged at 300 x g and 4°C for 20 min. The supernatant was collected, heat-inactivated at 56°C for 30 min and stored at −20°C until use. Sample analysis and cellular stimuli were performed with the pooled samples from the four groups mentioned.

**Table 1:** Plasma Donors.

| Group | Age (years) | Systolic blood pressure | Diastolic blood pressure | Height (m) | Weight (kg) | BMI (kg/m<sup>2</sup>) | Ethnicity | Bariatric surgery | Physical activity | Eating habits | Tobacco use | Alcohol use | STDs | Total hysterectomy | Hepatitis B (HBsAg) |
| --- | --- | --- | --- | --- | --- | --- | --- | --- | --- | --- | --- | --- | --- | --- | --- |
| Lean women | 43 | 118 | 73 | 150 | 45 | 20 | Mixed | No | Sedentary | Diet high in vegetables, fiber and white meat | No | Drinks socially | No | No | No |
| Lean women | 29 | 110 | 70 | 168 | 60 | 21 | Caucasian | No | Walking (>5 hours/week) | Diet high in red meat and fat, few vegetables | No | No |  | Yes | No |
| Lean women | 33 |  |  | 165 | 62 | 23 | Mixed | No | Sedentary | Diet high in vegetables, fiber and white meat | No | No | No | No |  |
| Lean women | 36 | 148 | 88 | 170 | 70 | 24 | Mixed | No | Sedentary | Diet high in vegetables, fiber and white meat | No | No | No | No | No |
| Averages Lean Women | 35.2 ± 3.0 | 125 ± 10 | 77 ± 5 | 163 ± 5 | 59 ± 5 | 22.0 ± 0.9 |  |  |  |  |  |  |  |  |  |
| Obese women | 38 | 142 | 78 | 168 | 75 | 27 | Caucasian | No | Walking (>2.5 hours/week) | Diet high in red meat and fat, few vegetable | No | No | No | No | No |
| Obese women | 34 |  |  | 162 | 70 | 27 | Not informed | No | Walking (>2.5 hours/week) | Diet high in vegetables, fiber and white meat | No | No | No | No | No |
| Obese women | 39 |  |  | 157 | 78 | 32 | Mixed | No | Sedentary | Diet high in red meat and fat, few vegetables | No | No | No | No | No |
| Obese women | 40 |  |  | 156 | 80 | 33 | Mixed | No | Sedentary | Diet high in vegetables, fiber and white meat | No | No | No | No | No |
| Obese women | 30 |  |  | 161 | 89 | 34 | Caucasian | No | Sedentary | Diet high in red meat and fat, few vegetables | No | Drinks socially | No | No | Yes |
| Obese women | 36 | 140 | 90 | 156 | 86 | 35 | Caucasian | No | Sedentary | Diet high in red meat and fat, few vegetables | No | No | No | No | No |
| Averages Obese Women | 36.1 ± 1.5 | 141 ± 1 | 84 ± 3 | 160 ± 2 | 80 ± 3 | 31.3 ± 1.4 |  |  |  |  |  |  |  |  |  |
| Lean men | 35 |  |  | 172 | 64 | 22 | Mixed | No | Sedentary | Diet high in vegetables, fiber and white meat | No | Drinks socially | No | Not applicable | No |
| Lean men | 37 | 120 | 62 | 170 | 65 | 22 | Mixed | No | Sedentary | Diet high in vegetables, fiber and white meat | No | No | No | Not applicable | No |
| Lean men | 32 | 124 | 79 | 184 | 78 | 23 | Mixed | No | Walking (>5 hours/ week) | Diet high in vegetables, fiber and white meat | No | Drinks socially | Yes | Not applicable | No |
| Lean men | 37 | 110 | 70 | 179 | 74 | 23 | Caucasian | No | Walking (>5 hours/ week) | Diet high in red meat and fat, few vegetables | No | Drinks socially | No | Not applicable | No |
| Lean men | 38 | 113 | 65 | 184 | 79 | 23 | Mixed | No | Walking (>2.5 hours/ week) | Diet high in vegetables, fiber and white meat | No | Drinks socially | No | Not applicable | No |
| Lean men | 32 |  |  | 170 | 68 | 24 | Black | No | Sedentary | Diet high in red meat and fat, few vegetables | No | No | Yes | Not applicable | No |
| Lean men | 36 |  |  | 174 | 74 | 24 | Caucasian | No | Walking (>5 hours/ week) | Diet high in red meat and fat, few vegetables | No | Drinks socially |  | Not applicable | No |
| Lean men | 32 |  |  | 180 | 81 | 25 | Caucasian | No | Walking (>5 hours/ week) | Diet high in red meat and fat, few vegetables | No | Drinks socially | No | Not applicable |  |
| Averages Lean Men | 35 ± 1 | 117 ± 1 | 69 ± 3 | 177 ± 2 | 73 ± 2 | 23.3 ± 0.4 |  |  |  |  |  |  |  |  |  |
| Obese men | 38 | 110 | 70 | 184 | 91 | 27 | Mixed | No | Sedentary | Diet high in red meat and fat, few vegetables | No | No | Yes | Not applicable | No |
| Obese men | 34 | 116 | 73 | 178 | 87 | 27 | Mixed | No | Sedentary | Diet high in vegetables, fiber and white meat | No | Drinks socially | No | Not applicable | No |
| Obese men | 37 | 110 | 85 | 170 | 80 | 28 | Caucasian | No | Walking (>5 hours/ week) | Diet high in red meat and fat, few vegetables | No | Drinks socially | No | Not applicable | No |
| Obese men | 40 |  |  | 170 | 80 | 28 | Caucasian | No | Sedentary | Diet high in vegetables, fiber and white meat | No | No | No | Not applicable | No |
| Obese men | 38 | 117 | 80 | 174 | 92 | 30 | Caucasian | No | Sedentary | Diet high in red meat and fat, few vegetables | No | No | No | Not applicable | No |
| Obese men | 35 |  |  | 165 | 83 | 30 | Caucasian | No | Sedentary | Diet high in red meat and fat, few vegetables | No | No | No | Not applicable | No |
| **Obese men** | 34 |  |  | 175 | 94 | 31 | Mixed | No | Walking (>2.5 hours/ week) | Diet high in vegetables, fiber and white meat | No | Drinks socially | No | Not applicable | No |
| **Obese men** | 34 |  |  | 180 | 115 | 35 | Mixed | No | Walking (>5 hours/ week) | Diet high in vegetables, fiber and white meat |  | Drinks socially | Yes | Not applicable | No |
| **Obese men** | 35 |  |  | 187 | 130 | 37 | Not informed | No | Sedentary | Diet high in red meat and fat, few vegetables | No | No | No | Not applicable | No |
| **Obese men** | 34 | 137 | 108 | 171 | 113 | 39 | Caucasian | No | Sedentary | Diet high in red meat and fat, few vegetables | No | No | No | Not applicable | No |
| **Averages Obese Men** | **36 ± 1** | **118 ± 4** | **83 ± 5** | **175 ± 2** | **97 ± 5** | **31.2 ± 1.4** |  |  |  |  |  |  |  |  |  |

### Cell cultures and incubations

INS-1E cells (a rat insulin-secreting β-cell line) were cultured with 100 IU/mL penicillin/streptomycin in RPMI-1640 medium (11.1 mM glucose, 10% bovine serum, 1 mM pyruvate, 10 mM HEPES, 2 mM glutamine and 0.1% β-mercaptoethanol) at 37°C and 5% CO_2_. Plating was done at 60,000 cells for all experiments. After 24 h, media was substituted for RPMI-1640 containing 10% fetal bovine serum (FBS, Gibco Standard Quality FBS, 12657-029, Made in Brazil), 10% serum from AL or CR animals; or 10% inactivated plasma from human volunteers. All experiments were performed 24 h after this medium change. In adiponectin supplementation experiments, cells were incubated with medium containing 10 µg/mL recombinant human adiponectin (SRP4901, Sigma-Aldrich, in sterile water), a quantity equivalent to typical adiponectin levels in lean women (Ding et al., 2012), in the absence or presence of other serum or plasma, as indicated.

### Cellular oxygen consumption

On the day of the experiment, cells were incubated in 500 μL RPMI-1640 without HEPES or FBS, containing 11.1 mM glucose for 1 h at 37°C, without CO_2_. During these 60 min, ports of the Seahorse cartridge containing oxygen probes were loaded with compounds to be injected during the assay (75 μL/port). The 24-well plate was then introduced into a Seahorse Bioscience XF24 analyzer (Billerica, MA, USA). Oxygen consumption was recorded for 30 min, at 5 min intervals, until system stabilization. Oligomycin was then injected at a final concentration of 1 µM, followed by carbonyl cyanide-4-(trifluoromethoxy) phenylhydrazone (FCCP) used at a final concentration of 10 µM, and antimycin (AA) and rotenone, both used at final concentrations of 2 μM. All respiratory modulators were used at optimal titrated concentrations, determined in preliminary experiments.

### Western blots

Serum and plasma samples were diluted in Laemmli buffer at a concentration of 1 μg/μL (human samples) or 7 μg/μL (animal samples). Particulate matter (see bellow) was diluted in PBS at the same volume as the original centrifugate and applied. Proteins were separated using a 12% denaturing polyacrylamide gel, transferred to nitrocellulose membranes, and incubated with anti-adiponectin antibody diluted 1:1000 (ab22554, Abcam). Ponceau staining was used as a loading control to avoid use of specific proteins that can fluctuate with metabolic changes. Fluorescent Secondary Anti-Rabbit Antibody diluted 1:10000 was added to the membranes and bands were visualized using an Odissey infrared system. Bands were semi-quantified by densitometric analysis using ImageJ software.

### Serum particulate matter isolation

Serum particulate matter was isolated by differential centrifugation, consecutively removing cells, apoptotic bodies, and cell debris. First, sera were submitted to 300 g for 5 min. The supernatant was collected and centrifuged at 2000 g for 10 min. This supernatant was collected and a centrifugation step at 10000 g for 1 h was performed. The resulting supernatant was submitted to a high-speed ultracentrifugation at 100000 g for 2 h to precipitate serum particulate matter. The resulting pellet was resuspended in phosphate buffered saline (PBS), at the same volume as the original serum sample, and stored at −80°C.

### Adiponectin immunoprecipitation

Rat sera were incubated at 4°C for 16 h in the presence of anti-adiponectin antibodies from Abcam (ab 56416) and Cell Signaling (2789s). Three different conditions were tested: Abcam antibodies at 1:10 dilution, Cell Signaling antibodies at 1:10, and a mixture of both antibodies at 1:20, as indicated in the figure legend. For the removal of antibody-adiponectin complexes or free antibodies, samples were incubated for 2 h at 4°C with protein A-Agarose (P2545) and then centrifuged for 1 min at 8000 x g. Prior to use, protein A-Agarose beads were washed and equilibrated in PBS. Adiponectin levels were evaluated by western blots, as described above.

### Cultured cell insulin secretion

Cells were plated as described above, and incubated with different serum or plasma samples 24 h later. After a further 24 h, they were pre-incubated for 30 min in Krebs-Henseleit (KH) solution containing 0.1% albumin and 5.6 mM glucose. They were then incubated for 1 hour at 37°C in the presence of 5.6 mM, 11.3 mM or 16.7 mM glucose. The supernatant was collected and stored at −20°C for subsequent measurements of secreted insulin. In addition, cells were lysed with acid-alcohol solution (52 mL ethanol, 17 mL water, 1 mL hydrochloric acid) to disrupt the cells and collect the intracellular insulin content. The determination of the amounts of secreted and intracellular insulin was performed following the Elisa Insulin Quantitation Kit protocol (Milipore, Billerica, MA, USA). Glucose-stimulated insulin secretion (GSIS) was calculated by dividing the concentration in the supernatant (secreted) by the remaining intracellular insulin (content).

### Pancreatic islet isolation and insulin secretion

Male Wistar rats (10 to 12 weeks) were deeply anesthetized with ketamine and xylazine, followed by decapitation. The abdomen was dissected, and the pancreas was inflated with 20 mL collagenase type V (0.7 mg/mL) in KH buffer. After full inflation, pancreases were removed and incubated for 25 min at 37°C, shaken manually, washed with KH buffer, and centrifuged three times at 1,000 rpm for 5 min. Islets were collected with a micropipette under a stereomicroscope and cultured for 24 h in RPMI-1640 medium containing 10 mM glucose, 1% penicillin/streptomycin and 10% FBS before receiving treatments. After 24 h, islets were randomly divided into wells under different conditions in 10 mM glucose RPMI: without FBS, with FBS, with 10 µg/mL adiponectin, or with FBS + adiponectin. After the 24 h incubation, the media were collected to check for insulin release over 24 hours, as well as LDH release (see below).

Islets were also checked for acute GSIS. Batches of 5 islets were collected in fresh tubes containing KH buffer with 5.6 mM glucose and incubated at 37°C for 30 min for stabilization. Supernatants were discarded and replaced by KH buffer with low (5.6 mM) or high (16.7 mM) concentrations of glucose and incubated at 37°C for 60 min. Insulin release in the medium over 24 hours and in the supernatant after acute stimulation with glucose was measured blindly by the Provet Institute (São Paulo, Brazil), by radioimmunoassay. Insulin concentrations are expressed as ng/mL.

### LDH quantification

Cultured INS-1E cells were plated and after 24 h treated with 10% plasma from lean and obese men and women for an additional 24 h, with or without 10 µg/mL of recombinant human adiponectin. Islets were incubated in 10 mM glucose for 24 h, with or without FBS and adiponectin, as described above. The culture medium supernatant was collected for lactate dehydrogenase (LDH) quantification, an enzyme that is released into the medium when there is damage to the plasma membrane. LDH activity was measured colorimetrically measuring NADH absorbance, following the protocol of a commercial quantification kit (Labtest, Lagoa Santa, MG, Brazil).

### Dada analysis

GraphPad Prism 7 was used for statistical evaluations. Data were expressed as means ± standard error of the mean (SEM) and statistically analyzed by unpaired Student’s t-tests or one-way ANOVA tests, with Tukey posttests. The minimum limit of significance was p < 0.05.

## Results

### Circulating factors induced by calorie restriction in rats protect β-cells in vivo and in vitro

In order to investigate the effects of circulating factors in the sera from lean versus mildly obese rats, we established a colony of animals in which a 60% caloric restriction (CR) diet, enriched with micronutrients to avoid malnutrition (Cerqueira et al., 2010), was introduced in early adulthood. These animals were compared to *ad libitum* (AL)-fed animals, which develop obesity, insulin resistance and other characteristics of the metabolic syndrome, and present significantly lower lifespans (Martin et al., 2010). Fig. 1A shows that the animals on the CR diet gained significantly less over the course of 15 weeks, but did not lose mass (which, if present, could be indicative of malnutrition). At the end of the intervention, the pancreases from both groups were collected, stained, and their islets quantified. We observed that pancreas from CR rats displayed an increased percentage of large islets compared to AL (Fig. 1B, p = 0.024), without changes in total islet area and average circularity (not shown). This is a moderate change in islet morphological distribution, compatible with the fact that the animals were obese, but not yet overtly diabetic.

**Figure 1 –.**
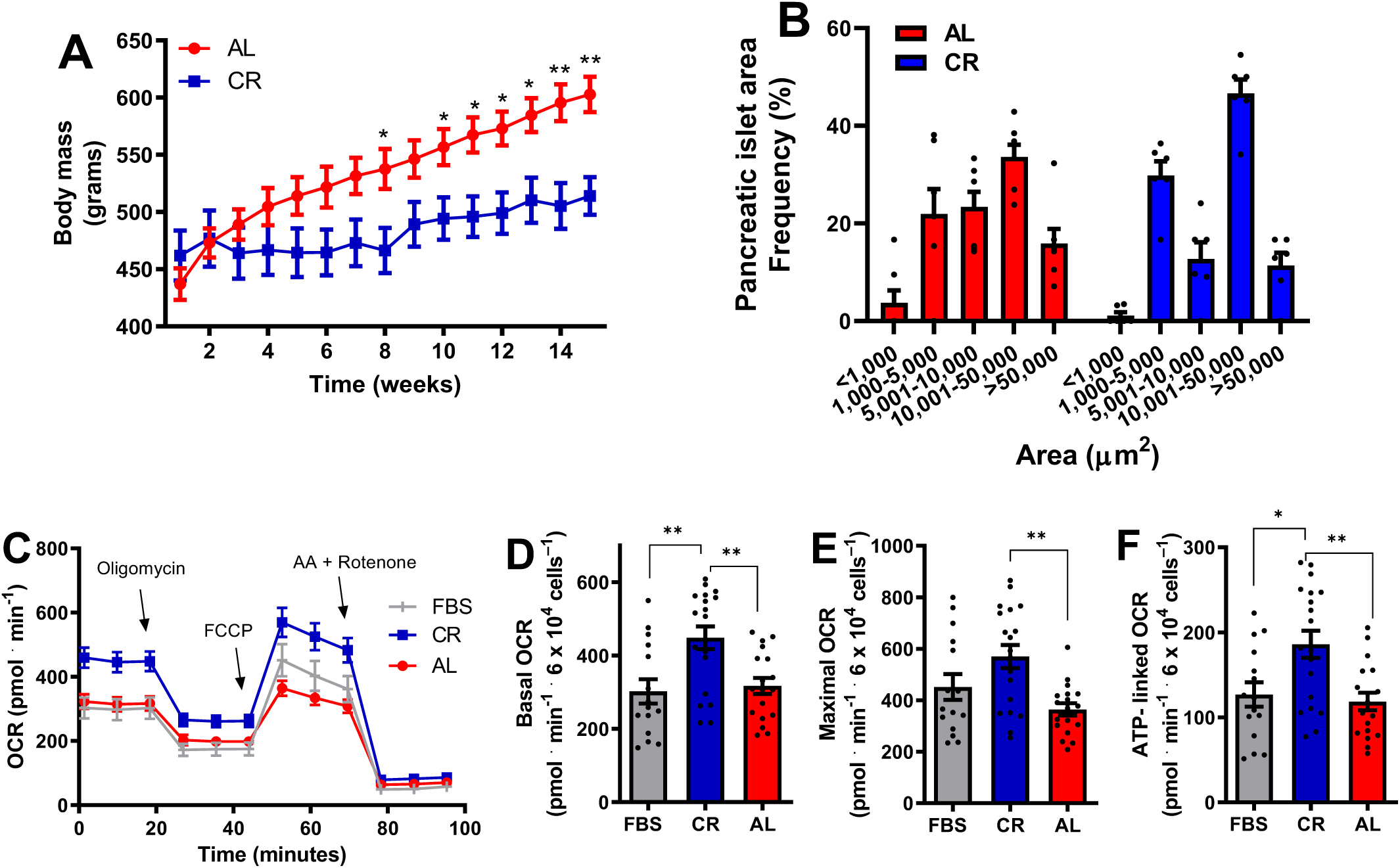
Sera from lean rats increases β-cell mitochondrial oxygen consumption. **A.** Weight progression of eight-week-old male rats on CR (blue) or AL (red) diets; *p < 0.05, **p < 0.005 compared to AL, unpaired t-student test, n = 6. **B.** Frequency (%) of islets from AL and CR rats versus area, in µm^2^. **C.** Typical INS1-E cell oxygen consumption rate (OCR) traces. Cells were incubated in media in the presence of the sera indicated, with 11.1 mM glucose. 1 µM oligomycin, 10 µM carbonyl cyanide-4-(trifluoromethoxy) phenylhydrazone (FCCP), and 2 µM antimycin plus rotenone were added when indicated. **D.** Basal OCR, quantified as initial minus antimycin plus rotenone-insensitive respiration, from traces such as those in panel B. **E.** Maximal OCR, quantified as FCCP-stimulated minus antimycin plus rotenone-insensitive respiration. **F.** ATP-linked OCR, quantified as initial minus oligomycin-insensitive respiration. Results are presented as means ± SEM, with individual dots representative of biological replicates. *p < 0.05, **p < 0.005, as indicated by one-way ANOVA with Tukey posttest; n = 15-19.

Next, sera from both groups were collected to be used on cultured INS-1E β-cells. As β-cell maintenance and its role in insulin secretion are dependent on oxidative phosphorylation, we tested the effects of culture media containing 10% serum from CR versus AL animals on oxygen consumption rates (OCR) in intact β-cells (Fig. 1C) by extracellular flux analysis. We found that 24 hours incubation with sera from lean CR animals results in higher OCRs relative to sera from obese AL animals (Cerqueira et al., 2016), and also results in higher OCRs relative to commercial fetal bovine serum (FBS). Enhanced OCRs were observed under basal conditions (Fig. 1D), which reflect normal mitochondrial oxygen consumption before the addition of any modulator (in traces such as shown in Fig. 1C), as well as conditions in which OCRs were maximized by uncoupler FCCP (Fig. 1E), which reflect the upper limits of mitochondrial electron transport chain capacity. CR sera also increased ATP-linked OCRs, which represent the difference between oxygen consumption under basal conditions and in the presence of ATP synthase inhibitor oligomycin (Fig. 1F), and are directly associated with ATP production and insulin secretion in these cells (Molina et al., 2009; Cerqueira et al., 2016). Overall, these findings confirm that sera from lean animals enhance oxidative phosphorylation activities in β-cells, known to be linked to glucose-stimulated insulin secretion.

### Circulating factors in human plasma regulate β-cell metabolic fluxes

As the effects of sera from lean animals are seen with 24 h incubations, they suggest the exciting idea that β-cell function, and hence their dysfunction in type 2 diabetes, can be acutely altered by the presence of circulating factors in the blood. Given the importance of this concept, we evaluated if the same was seen with human circulating factors. Human plasma samples stored in the Biobank of the *A. C. Camargo Cancer Center* were used. Although differences in clotting factors and nutrients exist between plasma and serum samples (Liu et al., 2010; Xu et al., 2011; Vignoli et al., 2022), human plasma samples from healthy donors were readily available in the Biobank, concurrently with extensive metabolic and nutritional information (Table 1), allowing for directed sample selection. Importantly, circulating hormonal factors do not differ much between plasma and serum samples (see, for example, Ashworth et al., 2021). Selected donors (Table 1) did not present chronic health conditions, smoke, or drink alcohol. They were within the same age range, but were clearly distinct in body mass indexes (BMI), which separated them into lean and obese groups: lean women (BMI 22 ± 0.9), obese women (BMI 31.0 ± 1.4), lean men (BMI 23.2 ± 0.3), and obese men (BMI 31.2 ± 1.3).

We then investigated the effects of diluted inactivated human plasma on β-cell metabolic fluxes in Fig. 2A. Typical OCR traces (Fig. 2A) were conducted under the same conditions as Fig. 1, and basal (Fig. 2B), maximal (Fig. 2C), and ATP-linked (Fig. 2D) OCRs were calculated. We found that samples incubated in the presence of plasma from lean women presented metabolic fluxes similar to those incubated in commercial FBS. On the other hand, plasma from obese women significantly suppressed basal, maximal, and ATP-linked OCRs. Plasma samples from lean men resulted in lower oxygen consumption compared to women’s samples, and OCRs were further suppressed in cells incubated with plasma from obese men. Overall, these results show a clear modulatory effect of circulating blood factors on metabolic fluxes in β-cells, which are stimulated by factors present in samples from lean and female subjects.

**Figure 2 –.**
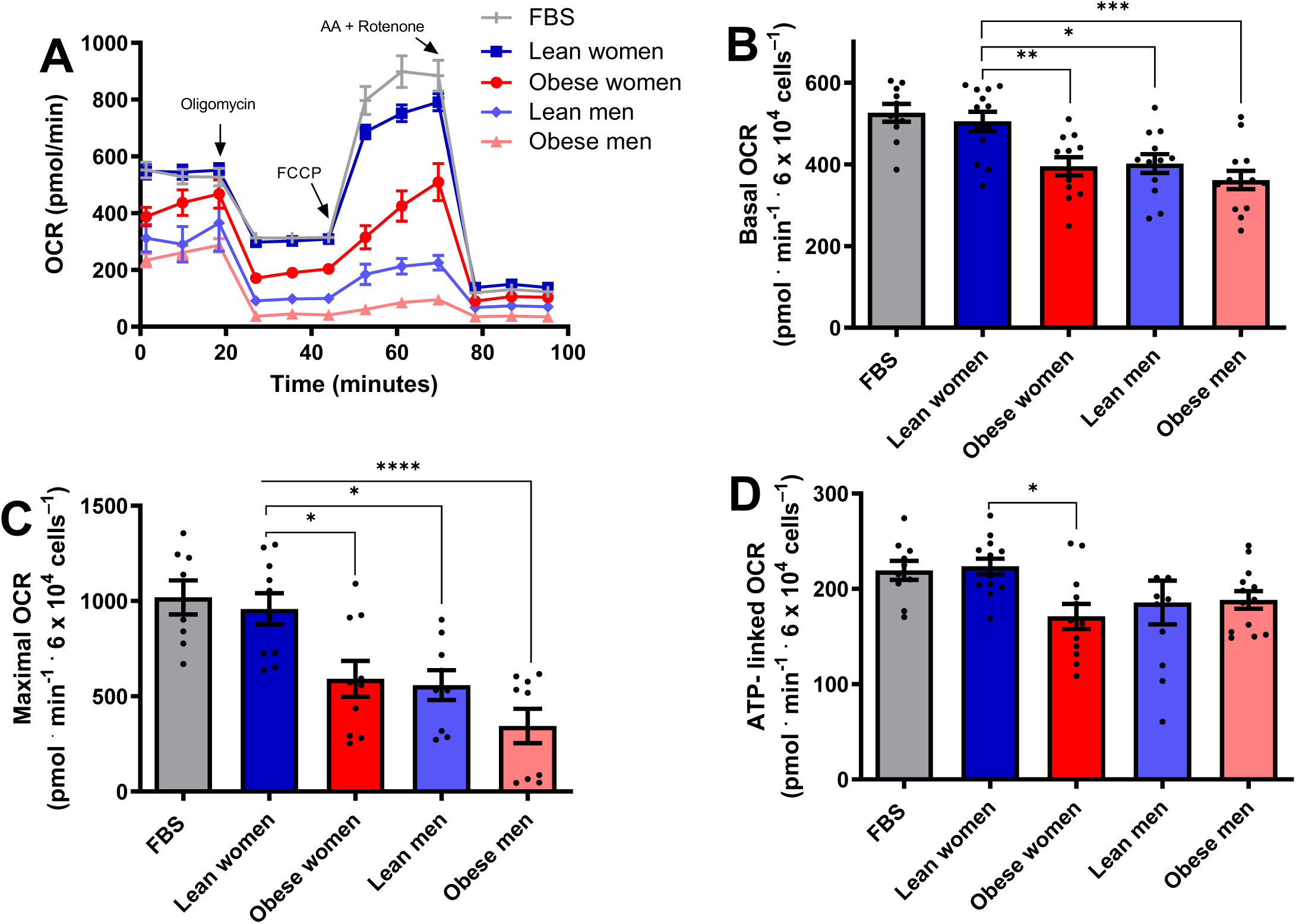
Female lean donor plasma increases β-cell mitochondrial oxygen consumption. **A.** Typical OCR traces, conducted under the same conditions as Fig. 1, with plasma samples from male, female, lean, and obese donors, as indicated. **B-D.** Basal, maximal and ATP-linked OCR, calculated as described in Fig.1. Results are presented as means ± SEM. *p <0.05, **p < 0.005***, p < 0.0005, ****p < 0.00005, one-way ANOVA test, with Tukey posttest, n = 8-13.

### Adiponectin, a circulating factor that modulates β-cell metabolic fluxes

We were interested in identifying the circulating blood factors responsible for these robust effects on β-cell metabolic responses. In peripheral tissues and vascular cells, enhanced mitochondrial electron transport capacity promoted by CR has been linked to adiponectin-activated eNOS signaling (Cerqueira et al., 2012). Indeed, adiponectin is an adipokine that modulates various metabolic processes, and is secreted by the adipose tissue in a manner increased by low body weights (Yang et al., 2001) and decreased by central adiposity (Arita et al., 1999). We thus quantified adiponectin in our rat serum samples (Fig. 3A), and found that the hormone was significantly increased in CR serum compared to AL. Indeed, prior work (Ding et al., 2012) has shown that animals under caloric restriction show a two-fold increase in circulating adiponectin levels. We also quantified adiponectin in human plasma (Fig. 3B) and found that both male and obese donors had decreased adiponectin, also consistent with prior work (Arita et al., 1999; Chandran et al., 2003). The levels of adiponectin in the blood therefore closely mirror the metabolic flux effects we observed in β-cells (Figs. 1 and 2).

**Figure 3 –.**
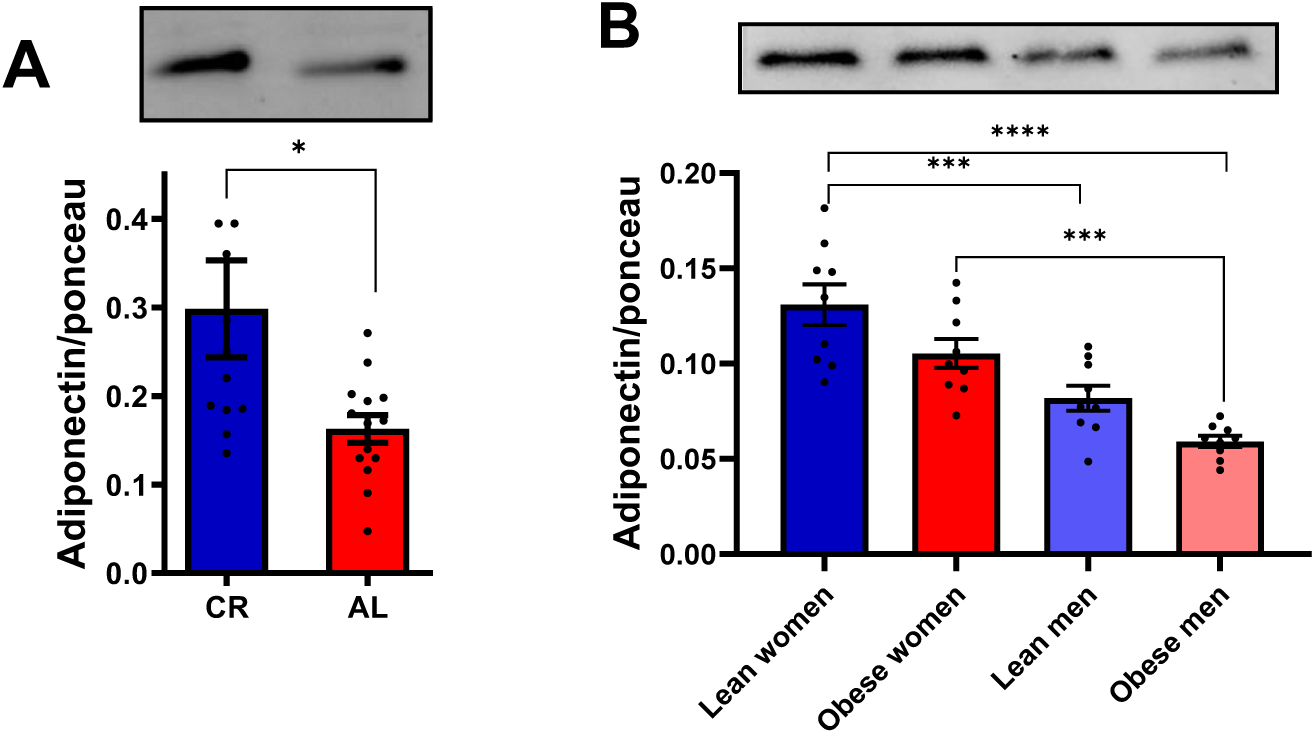
Biological sex and nutritional status alter the amount of circulating adiponectin. **A.** Quantification of adiponectin in serum samples from AL and CR rats; means ± SEM, *p < 0.05, unpaired t-student test, n = 12-14. **B.** Quantification of adiponectin in plasma samples from lean or obese men and women; means ± SEM; ***p < 0.0005, ****p < 0.00005, one-way ANOVA test with Tukey posttest, n = 9.

To investigate if adiponectin was responsible for the enhanced metabolic fluxes in β-cells promoted by lean sera, we attempted to remove the hormone from CR sera by immunoprecipitation, but found that metabolic fluxes were unaltered relative to full CR sera (Fig. 4A-D). We also found that adiponectin was very hard to fully deplete from the sera using different commercial antibodies at high concentrations, alone or in combination (Fig. 4E). Indeed, adiponectin is present in the circulation at very high concentrations relative to other adipokines and hormones. Additionally, it is present as trimers, hexamers, and high molecular weight multimers (Pajvani et al., 2003; Tsao et al., 2003; Waki et al., 2003); the latter form is the more active in improving insulin sensitivity and protecting against glucose intolerance (Hara et al., 2007). High molecular weight adiponectin (but not other adipokines such as leptin) is present in exosomes in significant quantities (Phoonsawat et al., 2014), which may hamper immunodepleting protocols. Consistently, we found that particulate matter (PM) from rat sera, obtained by ultracentrifugation, contained significant amounts of adiponectin (Fig 4E). Interestingly, while incubating β-cells under serum-free conditions hampers metabolic fluxes (Fig. 4F, black symbols), the addition of PM matter only from either commercial or rat sera was sufficient to re-establish much more robust oxygen consumption, suggesting membrane-associated serological components such as adiponectin are sufficient to promote β-cell metabolic fluxes.

**Figure 4 –.**
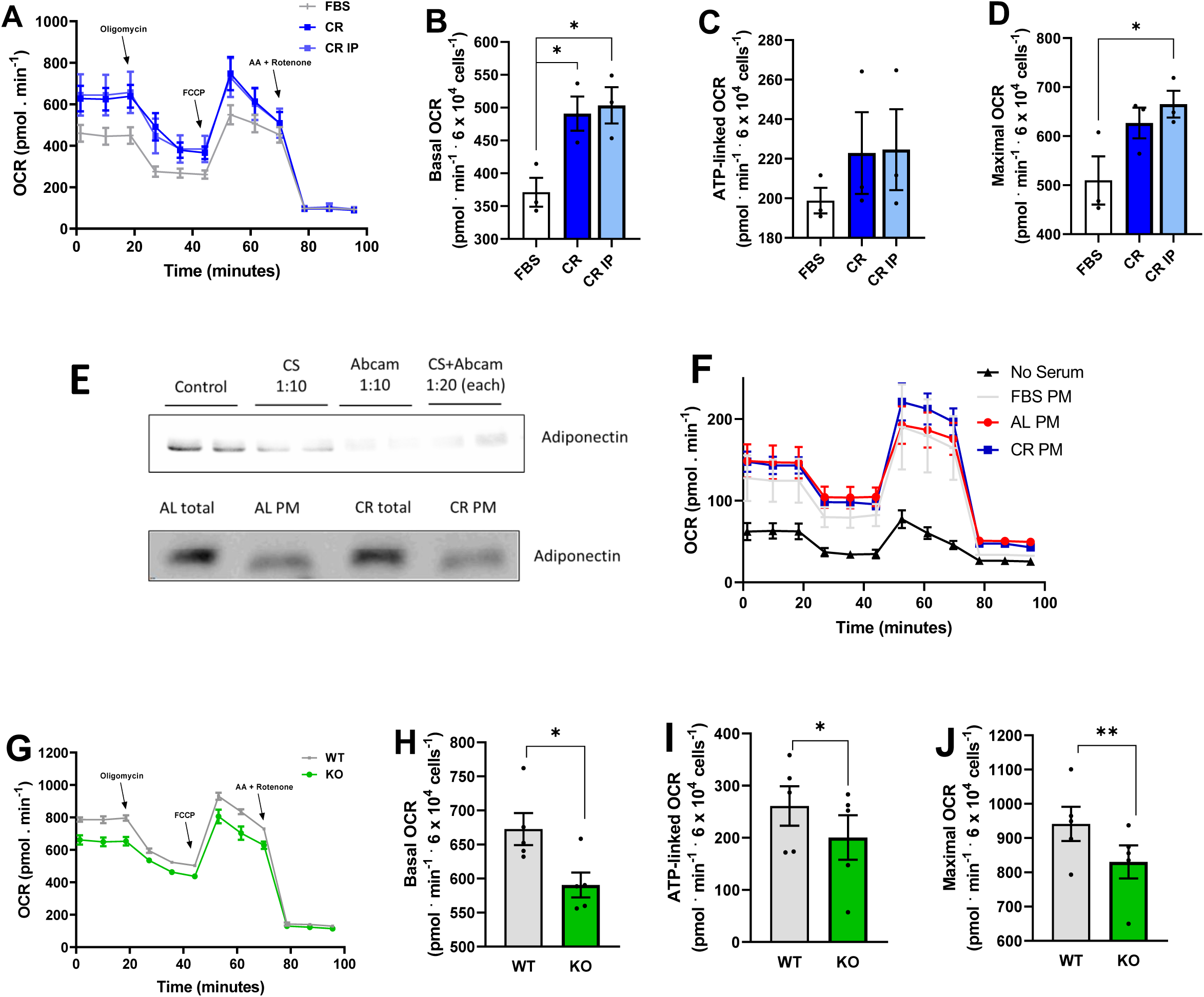
Serum adiponectin is partially membrane-bound and regulates metabolic fluxes. **A-D.** OCR traces and quantifications, conducted under the same conditions as Fig. 1, with commercial serum or CR rat serum depleted of adiponectin by immunoprecipitation (IP) with Abcam antibodies, as indicated. *p<0.05, one-way ANOVA, n = 3. **E.** Upper panel: Typical western blot showing that adiponectin is not completely removed from rat sera through immunoprecipitation with Cell Signaling (CS) and/or Abcam antibodies. Primary antibodies were tested alone (dilution 1:10) and in combination (dilution 1:20). Lower panel: Typical western blot showing adiponectin in full sera and the particulate matter (PM) from AL or CR animals. **F.** OCR measurements under conditions similar to Fig. 1, in the absence of serum or the presence of PM from FBS, CR or AL sera suspended in PBS. **G-J.** OCR traces and quantifications, conducted under the same conditions as Fig. 1, with full sera from WT mice or adiponectin KO animals. *p < 0.05, **p < 0.005, Student’s t test, n = 5.

In order to verify if adiponectin was an important component without the ability to remove it due to its binding to membranes, we treated the cells with sera from WT mice or their KO counterparts unable to produce adiponectin (Fig. 4G-J). While oxygen consumption was present in cells incubated with KO sera, it was significantly lower than in WT sera under all respiratory conditions. These data indicate that circulating membrane-bound factors, including adiponectin, are regulators of β-cell metabolism.

Given the finding that PM matter from sera alone can support oxygen consumption in β-cells (Fig. 4F), we decided to investigate the direct effects of purified adiponectin on β-cell metabolic fluxes (Fig. 5A-D). While robust OCRs are present in FBS, oxygen consumption was strongly suppressed in the absence of any sera. Impressively, adding recombinant adiponectin alone, in the absence of serum, at quantities compatible with those present in CR sera (10 µg/mL, Ding et al., 2012), promoted metabolic fluxes close to those seen in the presence of commercial FBS. This shows that adiponectin is sufficient to promote a strong and previously undescribed effect supporting oxidative phosphorylation in β-cells in the absence of any other serological factor.

**Figure 5 –.**
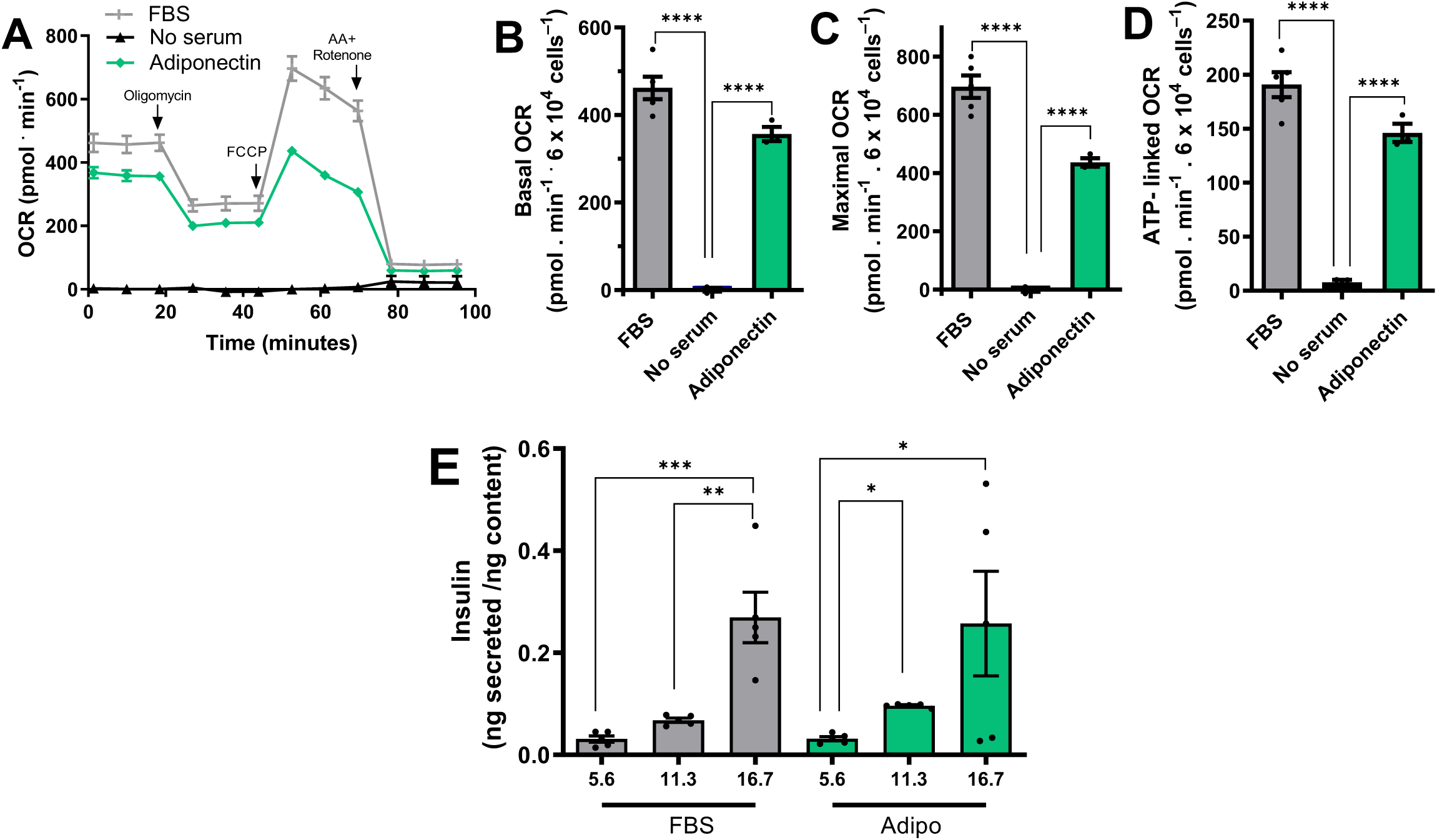
β-cell metabolic fluxes are strongly induced by adiponectin at levels present in lean sera. Experiments were conducted under similar conditions to Fig. 1, in the presence of FBS, no sera, and/or 10 µg/mL adiponectin, as indicated. **A.** Typical traces. **B-D.** Basal, maximal, and ATP-linked OCR. **E.** Insulin secretion corrected for cellular insulin content was measured after one hour of incubation with 5.6, 11.3, or 16.7 mM glucose, in the presence of FBS or adiponectin, as indicated. Results are presented as means ± SEM. *p <0.05, ****p < 0.00005, one-way ANOVA with Tukey posttest, n = 3-5.

Given the strong adiponectin effects on OCRs, we questioned if it could also promote glucose-stimulated insulin secretion (GSIS) in INS-1E β-cells, which are known to depend on the presence of serum (Arumugam et al., 2008). Fig. 5E compares insulin secretion with different glucose concentrations in cells incubated in FBS or adiponectin in the absence of FBS. Our data show that adiponectin alone was able to promote GSIS patterns similar to those seen in full serum. Indeed, responses were not significantly different from those seen in FBS controls. This demonstrates that adiponectińs metabolic effect increasing ATP-linked OCRs is accompanied by the expected increase in insulin release from β-cells. It also confirms that adiponectin has a pronounced impact on β cell function, in isolation.

The finding that adiponectin strongly protects cultured β-cells in the absence of any other serological factor is highly surprising, and prompted us to verify whether primary β-cells are also similarly responsive to adiponectin. Figures 6A and 6B show measurements of insulin release from primary rat islets in the presence of low and high glucose, respectively. As expected, insulin was secreted in high glucose (Fig. 6B) when FBS was present, but not in its absence. Strikingly, the presence of recombinant adiponectin alone restored islet secretory function, in a manner that was not additive to the presence of FBS (Fig. 6B), indicating that adiponectin itself is sufficient to promote β-cell-protective effects found with commercial serum.

**Figure 6 –.**
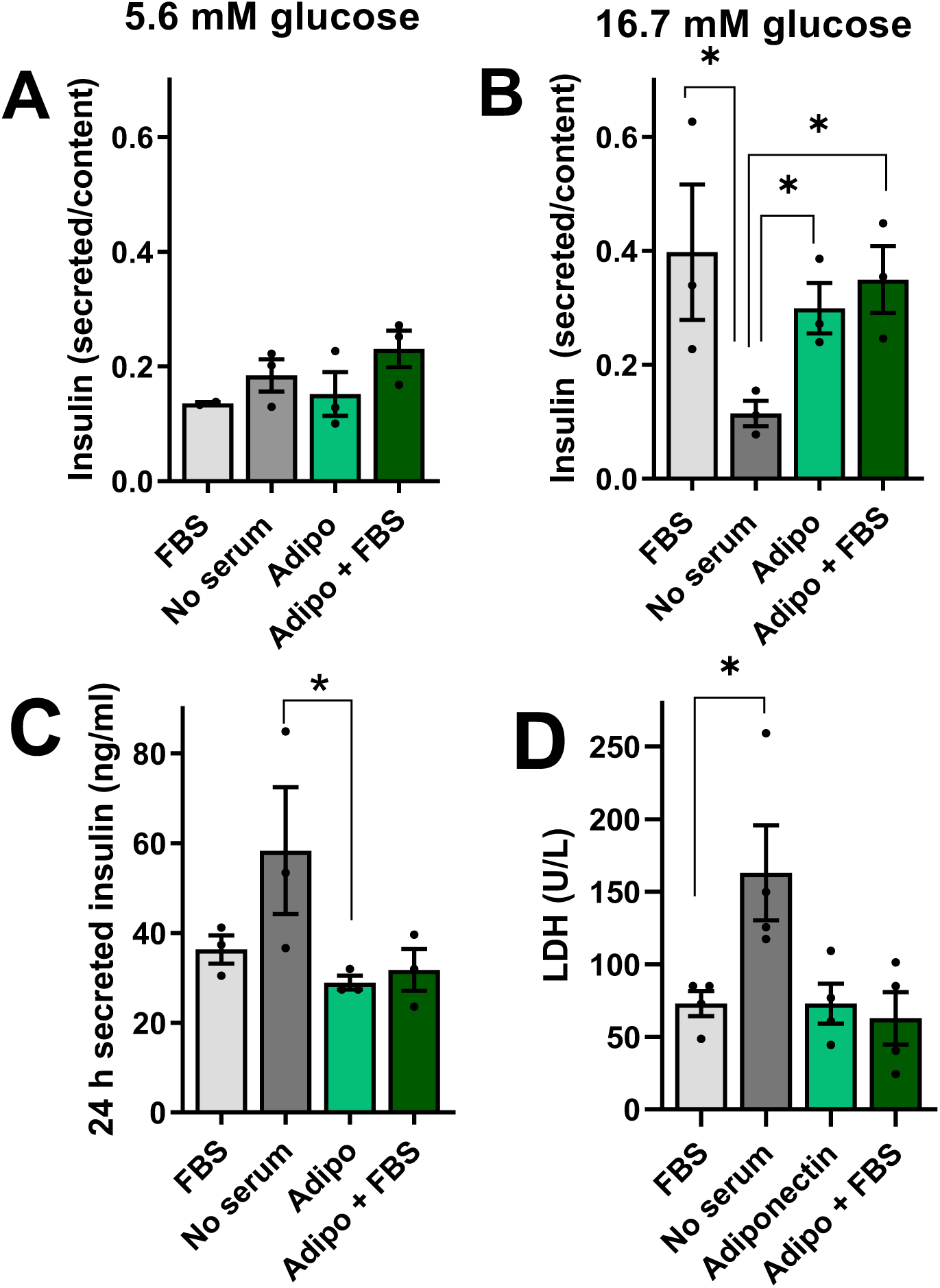
Serum adiponectin is necessary to maintain islet function and integrity. Islet insulin secretion was measured after 60 min as described in the Methods section with low (5.6 mM, Panel **A**) or high (16.7 mM, Panel **B**) glucose in the presence of FBS, no serum, or 10 µg/mL adiponectin, as indicated. **C.** Insulin secretion collected from islets incubated in 10 mM glucose over a 24 h period, as an indication of cell integrity loss. **D.** LDH activity measured in the culture media. *p <0.05, one-way ANOVA with Tukey posttest, n = 3-4.

The lack of insulin release in media with high glucose and no serum is a consequence of islet damage, as indicated by the fact that 24 h insulin secretion (due to β-cell membrane disruption, leading to intracellular insulin release independent of glucose stimulation, Fig. 6C) and LDH release (Fig. 6D) were augmented in islets incubated with no sera. Once again, adiponectin alone was protective against these insults, in a manner that was not additive with the effect of FBS, demonstrating that this hormone is strongly protective toward primary islets as well as cultured β-cell lines.

### Adiponectin supplementation reverses the damaging effects of obese plasma on β-cells

Next, we sought to verify if adiponectin supplementation in the plasma from obese donors was capable of reversing the metabolic flux limitations observed in Fig. 2. We monitored OCRs in samples in which 10 μg/mL adiponectin was added together with the plasma of obese men and women (Fig. 7). Once again, the effect of added adiponectin was very pronounced: it was able to stimulate OCRs under all conditions, in both male and female obese plasma-treated β-cells. These results suggest that samples from male and obese subjects do not primarily decrease metabolic fluxes in β-cells due to the presence of damaging circulating molecules, but instead as a result of the lack of the stimulatory factor adiponectin.

**Figure 7 –.**
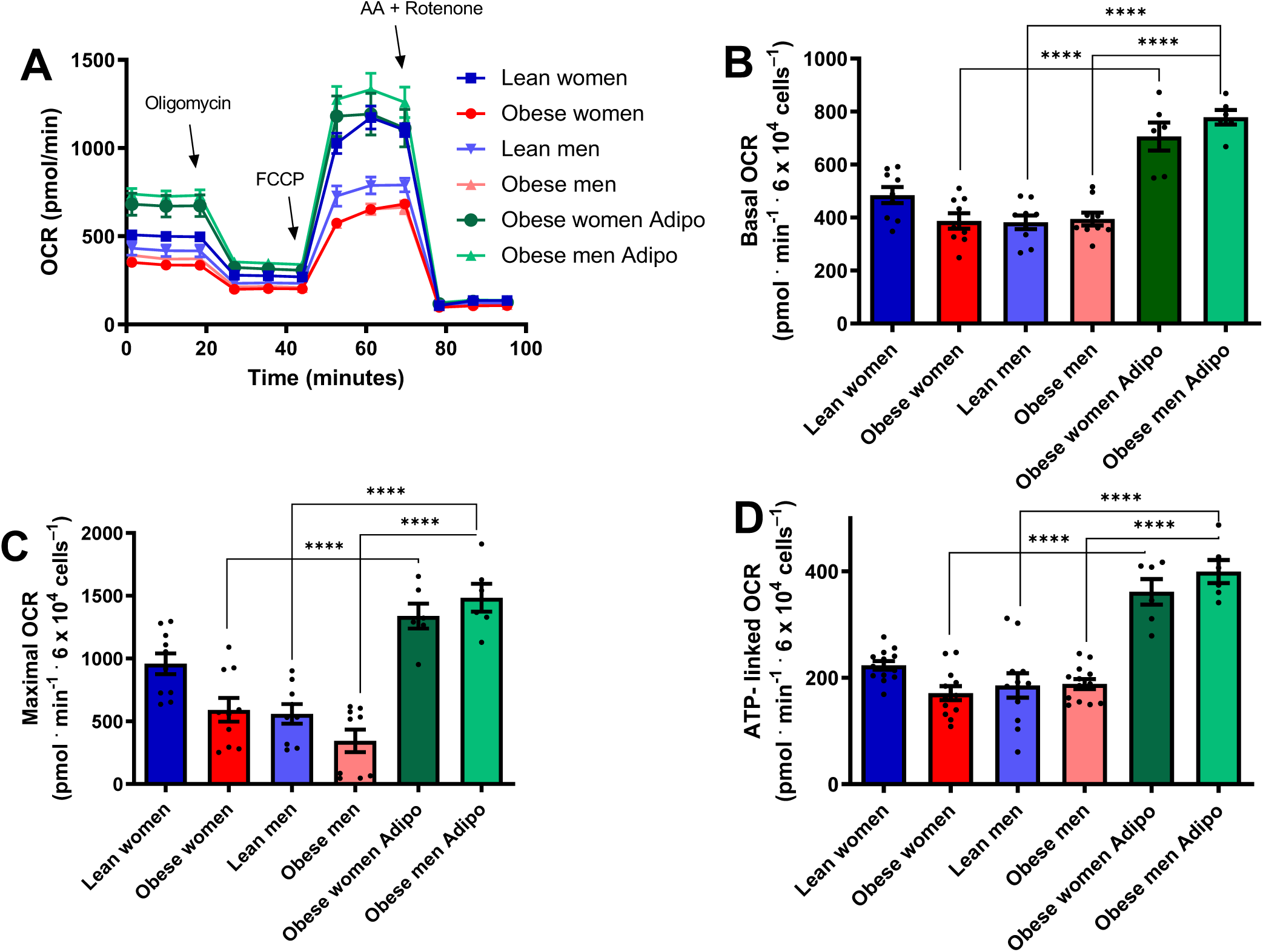
Low β-cell metabolic fluxes in obese and male sera are reversed by adiponectin. **A.** Typical OCR traces, conducted under the same conditions as Fig. 2, with plasma samples from male, female, lean, and obese donors, in the presence of 10 µg/mL adiponectin, where indicated. **B-D.** Basal, maximal, and ATP-linked OCR. Results are presented as means ± SEM. ****p < 0.00005, one-way ANOVA test, with Tukey posttest, n = 4-13.

Given the strong protective results on metabolic fluxes, we also tested the effect of added adiponectin on GSIS. In samples incubated for 24 h in plasma from lean women, an expected increase in insulin release was observed with raising glucose concentrations (Fig. 8A). Incubation in plasma from obese women lead to very high levels of insulin secretion, without any effect of glucose concentrations. This could be due to the presence of ruptured cells, releasing insulin into the supernatant. Indeed, when we quantified LDH release as a measure of cell integrity (Fig. 8B), we found that the sera of obese women promoted significant cell damage. Strikingly, the addition of adiponectin to the samples incubated with plasma of obese women completely prevented LDH release (Fig. 8B) and restored functional GSIS (Fig. 8A). Adiponectin also prevented LDH release from cells incubated in plasma from obese male donors (Fig. 8D), and restored expected insulin secretion patterns (Fig. 8C), which were not observed in the plasma from obese male donors alone. Overall, these results demonstrate that the presence of adiponectin is capable of completely abrogating the damaging effects of obese plasma on β-cells.

**Figure 8 –.**
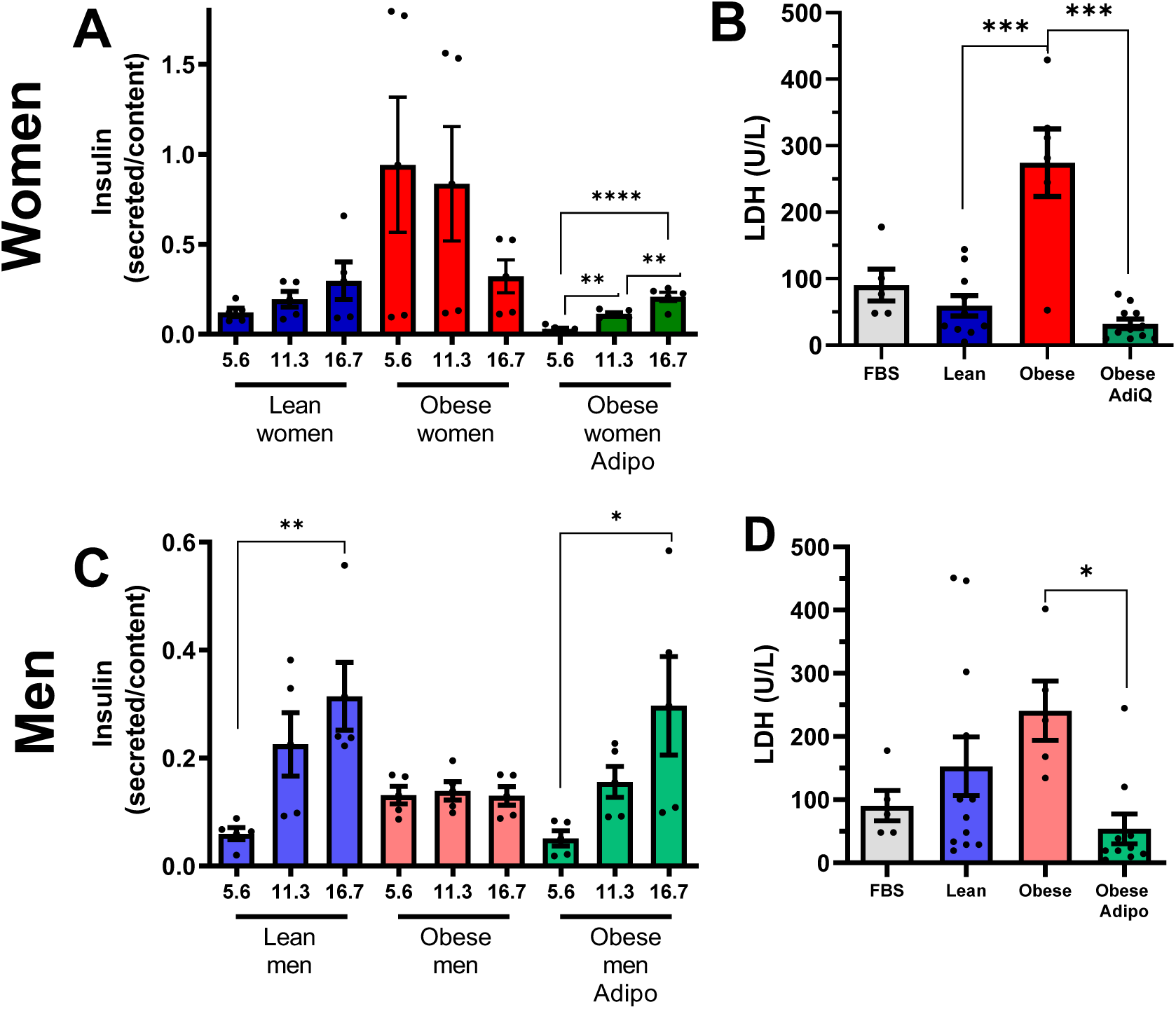
Adiponectin protects β-cell function and integrity against damage promoted by plasma from obese donors. **A, C.** Insulin secretion corrected for cellular insulin content after one hour of incubation with 5.6, 11.3, or 16.7 mM glucose, as indicated. Mean ± SEM, **p < 0.005, ***p < 0.0005, ****p < 0.00005, one-way ANOVA test with Tukey posttest, n = 5. **B, D.** Lactate dehydrogenase (LDH) in culture media after incubation with 10% FBS or plasma from lean or obese men and women, with or without adiponectin, as shown. Mean ± SEM, *p < 0.05, ***p < 0.0005, one-way ANOVA with Tukey posttest, n = 5-12.

## Discussion

Understanding mechanisms involved in interventions known to extend longevity, such as CR, is essential to uncover new target pathways to promote health aging. Previous studies by our group found that serum from CR rats diluted in culture media is capable of increasing the expression of proteins involved in mitochondrial fusion (mitofusin-2, Mfn-2; optic atrophy 1, OPA-1) and reducing DRP-1, involved in mitochondrial fission. This results in increased mitochondrial length and connectivity in β-cells, an effect that promotes glucose-stimulated ATP production and insulin secretion (Cerqueira et al., 2016). Sera from CR animals was thus protective against *in vitro* toxic stimuli (glucolipotoxicity with 20 mM glucose and 0.4 mM palmitate) that are a model for type 2 diabetes, preventing mitochondrial fragmentation and respiratory dysfunction, as well as preserving GSIS (Cerqueira et al., 2016). Knowing that the sera could not contribute significantly to the nutrient pool at the dilution used, we hypothesized that the results were due to the presence of hormonal circulatory components.

Other studies have tested the *in vitro* effects of sera from calorically-restricted animals. In normal human diploid fibroblasts, serum from CR animals delayed senescence and significantly increased longevity compared to serum from AL animals (de Cabo et al., 2015). In human hepatoma cells, serum from calorie-restricted volunteers increased longevity markers such as Sirtuin 1 and PGC-1a, and enhanced stress resistance (Allard et al., 2008). In vascular cells, rat CR sera activates the insulin pathway and phosphorylation of endothelial nitric oxide synthase (Cerqueira et al., 2012), which modulates mitochondrial biogenesis, a process believed to have a central role in the longevity effects of CR (Nisoli et al., 2005). The beneficial effects of CR sera on vascular cells were eliminated when the sera were depleted of adiponectin (Cerqueira et al., 2012), suggesting this is an important hormonal factor modulating oxidative phosphorylation in lean animals.

In the present study, we find that the beneficial effects of CR animal circulatory factors are also found using plasma from lean female donors. We also shed light on adiponectin, an adipokine enriched in lean females, as a protective component of sera from lean subjects on β-cells. Indeed, we found that sera from KO mice devoid of adiponectin decreases metabolic fluxes, while adiponectin itself (even in the absence of sera) is a strong determinant of β-cell metabolic fluxes and GSIS, in both cell lines and primary rat β-cells. Additionally, supplementation of this hormone reverses the damaging effects of obese human plasma on β-cells, suggesting that its low levels, rather than the presence of damaging signaling molecules, are determinant for β-cell damage.

Adiponectin has been extensively studied in aging and the metabolic syndrome, using both rodent models and populational observations. Interestingly, adiponectin levels do not decline with age - they increase in both men and women, apparently due to lower clearance by the kidneys (Isobe et al., 2005). Despite this, lifetime levels of this adipokine are closely related to aging, as knockout animals present lower longevity, while transgenic adiponectin overexpression prolonged median lifespan (Otabe et al., 2007; Li et al., 2021). Additionally, adiponectin KO mice have exacerbated age-related glucose and lipid metabolism disorders (Li et al., 2021). Conversely, PEGylated AdipoRon derivatives, capable of overcoming the poor bioavailability of these adiponectin receptor-activating molecules, improve glucose metabolism in mice fed high fat diets (Onodera et al., 2021). Consistently, circulating adiponectin is lower in individuals with type 2 diabetes compared to healthy individuals, and well as those with risk factors such as obesity (Hotta et al., 2000; Xydakis et al., 2004; Bahia et al., 2006; Choi et al., 2007; Eglit et al., 2013). Furthermore, transgenic mice overexpressing human adiponectin and fed with a high-fat/high-sucrose diet present increased longevity as well as reduced morbidity and mortality (Otabe et al., 2007). These animals also showed reduced body weight, and less accumulation of subcutaneous and visceral fat, with smaller adipocytes in both tissues. This, added to an increase in oxygen consumption associated with equal caloric consumption compared to control animals, suggests an increase in energy expenditure (Otabe et al., 2007). On the other hand, adiponectin-deficient animals exhibit higher body mass, impaired glucose tolerance, and more triacylglycerol accumulation than control animals (Kubota et al., 2002; Maeda et al., 2002; Mawrocki et al., 2006; Streeper et al., 2006). In line with this, a meta-analysis of human populational studies shows a strong correlation between low adiponectin levels and type 2 diabetes risk, even when analyzed independently of ethnic background and inflammatory markers (Wang et al., 2018). This correlation is especially significant with high molecular weight complexes of this adipokine (Fisher et al., 2005; Schraw et al., 2008).

Indeed, adiponectin is found in the plasma as complexes (trimers, hexamers, high molecular weight multimers), that have distinct biochemical characteristics and physiological outcomes (Schraw et al., 2008). The fact that we found that recombinant adiponectin added to the plasma from obese humans can reverse the impairment of metabolic fluxes and insulin secretion indicates that specific forms of adiponectin aggregates are not the primary reason for the effect of obese plasma, but rather lack of total adiponectin. Unfortunately, because the sera and plasma in our studies are heat-inactivated to allow incubations with cells, we were unable to resolve these samples in native gels and confirm the changes in aggregation of adiponectin expected (protein aggregates unable to enter native gels were formed, results not shown). We were, however, able to identify that adiponectin was present in the particulate matter of our sera, and that this membrane-associated fraction was active in β-cells. This is compatible with the finding that exosomes contain significant amounts of high molecular weight, active, adiponectin (Phoonsawat et al., 2014).

Adiponectin acts on cells by binding to adiponectin receptors and modulating the phosphorylation of protein kinase B (AKT) (Cerqueira et al., 2012; Su et al., 2015; Xu et al., 2018). Specifically in pancreatic β-cells, adiponectin regulates insulin gene expression, in addition to increasing cell viability and decreasing apoptosis via AKT phosphorylation (Wijesekara et al., 2010), results compatible with the protective effects of adiponectin we observed in this study. Adiponectin also acts via the hypothalamus to inhibit appetite and increase energy expenditure (Qi et al., 2004; Yu et al., 2020), promoting fatty acid oxidation in muscle and liver, leading to weight loss (Fruebis et al., 2001; Yamaguchi et al., 2001). Overall, these studies indicate that this hormone is a major regulator of energy metabolism in various organs, and are in line with our added finding of its strong metabolic and functional effects directly on b-cells.

In addition to adiposity, biological sex also affects the amount of adiponectin, with more of this hormone circulating in women than in men (Nishizawa et al., 2002; Snijder et al., 2006; Rathmann et al., 2007; Ahonen et al., 2009; Eglit et al., 2013). The reduction in men appears to be linked to the presence of higher concentrations of testosterone, as this hormone reduced adiponectin concentrations in mice, and castration induced an increase in plasma adiponectin associated with a significant improvement in insulin sensitivity (Nishizawa et al., 2002). In fact, although obesity is more common in women, type 2 diabetes is more often diagnosed at a lower age and with lower body mass indexes in men (Kautzky-Willer et al., 2016; Peters et al., 2019; Saeedi et al., 2019). It is tempting to speculate that this may be related to the lack of adiponectin and its β-cell-protective effects, as uncovered in this study.

### Experimental limitations

Unfortunately, due to the large amount of serum samples needed, we were not able to conduct studies in animals and humans of different age groups. Also, we were unable to study the aggregation forms of adiponectin, as sera and plasma must be heat-inactivated prior to cell culture incubations, hampering resolution under native conditions. Finally, our studies involving added adiponectin used the recombinant protein at levels found in lean individuals, and thus added to adiponectin present, possibly explaining the overshoot of oxygen consumption rates above those seen with sera from lean subjects.

This study does not include widespread analysis of differences in the circulating factors in lean and obese rodents and humans in an attempt to uncover factors involved in the protection of β-cells. While this approach fails to uncover the full range of differences dietary interventions promote uncovered in multi-omic studies (Dao et al., 2019; Aon et al., 2020), which may include many other molecules with impact on β-cells such as inflammatory factors, it has the advantage of isolating and deepening the understanding of the specific effects of one adipokine, adiponectin, on β-cell function.

## Conclusions

We show here for the first time that adiponectin, at quantities present in the blood of lean long-lived rats and mice, as well as lean women donors, is a strong stimulatory factor for metabolic fluxes in β-cells. The presence of adiponectin alone, in the absence of any other serological factor, sustains ATP-linked respiration and associated glucose-stimulated insulin release in primary and cultured β-cells. Addition of adiponectin to plasma-supplemented media also rescues β-cell function compromised by incubation with samples from obese donors. Overall, our results suggest this hormone, its receptors, and the signaling pathways it activates are robust potential targets for treatment in obesity and aging-related disruption of glucose-stimulated insulin secretion.

## Conflict of interest

The authors declare no conflicts of interest.

## Funding

This study was funded by the Conselho Nacional de Desenvolvimento Científico e Tecnológico (CNPq), Coordenação de Aperfeiçoamento de Pessoal de Nível Superior (CAPES) Finance Code 001, and Fundação de Amparo à Pesquisa no Estado de São Paulo (FAPESP) grants 19/19644-1, 20/06970-5 and 13/07937-8, as part of the Centro de Pesquisa, Inovação e Difusão de Processos Redox em Biomedicina (Redoxoma). ACM, JDCS, and EVB are supported by FAPESP fellowships (19/04233-6, 19/05226-3, 21/02481-2). Biobanking of human samples was supported by the Programa Nacional de Apoio à Atenção Oncológica (PRONON), Banco de Tumores para Pesquisa em Tratamento, Prevenção e Diagnóstico Precoce do Câncer (25000.158.661/2014-68).

## Acknowledgments

We acknowledge Gisela Ramos Terçarioli and Cruz Alberto Mendoza Rigonati for histological technical assistance and Roberta Andrejew for experimental help in slide scanning. The authors also acknowledge to the A. C. Camargo Biobank for the plasma samples from normal donors. Community revision provided by AsapBio to the preprint of this article was greatly appreciated.

## Data sharing

Raw data will be provided upon request.

## Author contributions

ACM, JDCS, EAVB, and CCCS acquired experimental data. TGS and VMR participated in Biobank data and sample acquisition. FCM and SLF participated in WT and KO mouse sera acquisition. All authors participated in experimental design, analysis, and interpretation of data. All authors participated in drafting or revising the manuscript, and approved the final submitted version. All authors agree to be accountable for all aspects of the work, and in ensuring that questions related to the accuracy or integrity of any part of the work are appropriately investigated and resolved.

## Abbreviations

Antimycin A (AA); *ad libitum* (AL); adiponectin (Adipo); body mass index (BMI); calorie restricted (CR); carbonyl cyanide-4-(trifluoromethoxy) phenylhydrazone (FCCP); ethylenediaminetetraacetic acid dipotassium salt (EDTA); fetal bovine serum (FBS); glucose-stimulated insulin secretion (GSIS); immunoprecipitated (IP); Knockout (KO); lactate dehydrogenase (LDH); oxygen consumption rate (OCR); standard error of the mean (SEM); Wild-type (WT)

## Notes

### Competing Interest Statement

The authors have declared no competing interest.

### Summary of Updates

New experimental data and discussions.

